# Analysis of overlapping genetic association in type 1 and type 2 diabetes

**DOI:** 10.1101/2020.06.17.156778

**Authors:** Jamie R.J. Inshaw, Carlo Sidore, Francesco Cucca, M. Irina Stefana, Daniel J. M. Crouch, Mark I. McCarthy, Anubha Mahajan, John A. Todd

## Abstract

**Aims/hypothesis:** Given the potential shared aetiology between type 1 and type 2 diabetes, we aimed to identify any genetic regions associated with both diseases. For associations where there is a shared signal and the allele that increases risk to one disease also increases risk to the other, inference about shared aetiology could be made, with the potential to develop therapeutic strategies to treat or prevent both diseases simultaneously. Alternatively, if a genetic signal colocalises with divergent effect directions, it could provide valuable biological insight into how the association affects the two diseases differently.

**Methods:** Using publicly available type 2 diabetes summary statistics from a genomewide association study (GWAS) meta-analysis of European ancestry individuals (74,124 cases and 824,006 controls) and type 1 diabetes GWAS summary statistics from a meta-analysis of studies on individuals from the UK and Sardinia (7,467 cases and 10,218 controls), we identified all regions of 0.5 Mb that contained variants associated with both diseases (false discovery rate<0.01). In each region, we performed forward stepwise logistic regression to identify independent association signals, then examined colocalisation of each type 1 diabetes signal with each type 2 diabetes signal using *coloc*. Any association with a colocalisation posterior probability of ≥0.9 was considered a genuine shared association with both diseases.

**Results:** Of the 81 association signals from 42 genetic regions that showed association with both type 1 and type 2 diabetes, four association signals colocalised between both diseases (posterior probability ≥0.9): (i) chromosome 16q23.1, near Chymotripsinogen B1 (*CTRB1*) / Breast Cancer Anti-Estrogen Resistance Protein 1 (*BCAR1*), which has been previously identified; (ii) chromosome 11p15.5, near the Insulin (INS) gene; (iii) chromosome 4p16.3, near Transmembrane protein 129 (*TMEM129*), and (iv) chromosome 1p31.3, near Phosphoglucomutase 1 (*PGM1*). In each of these regions, the effect of genetic variants on type 1 diabetes was in the opposite direction to the effect on type 2 diabetes. Use of additional datasets also supported the previously identified colocalisation on chromosome 9p24.2, near the GLIS Family Zinc Finger Protein 3 (*GLIS3*) gene, in this case with a concordant direction of effect.

**Conclusions/interpretation:** That four of five association signals that colocalise between type 1 diabetes and type 2 diabetes are in opposite directions suggests a complex genetic relationship between the two diseases.

**Research in Context:** **What is already known about this subject?**

- Other than insulin, there are currently no treatments for both type 1 and type 2 diabetes.
- Findings that genetic variants near the GLIS3 gene increase risk of both type 1 and type 2 diabetes have indicated shared genetic mechanisms at the level of the pancreatic β cell.

**What is the key question?**

- By examining chromosome regions associated with both diseases, are there any more variants that affect risk of both diseases and could support common mechanisms and repositioning of therapeutics between the diseases?

**What are the new findings?**

- At current sample sizes, there is evidence that five genetic variants in different chromosome regions impact risk of developing both diseases.
- However, four of these variants have the opposite direction of effect in type 1 diabetes compared to type 2 diabetes, with only one, near *GLIS3*, having a concordant direction of effect.

**How might this impact on clinical practise in the foreseeable future?**

- Genetic findings have furthered research in type 1 and type 2 diabetes independently, and suggest therapeutic strategies. However, our current investigation into their shared genetics suggests that repositioning of current type 2 diabetes treatments into type 1 diabetes may not be straightforward.

## Introduction

There is a genetic component to both type 1 and type 2 diabetes, with approximately 60 chromosome regions associated with type 1 diabetes^1^ and over 200 associated with type 2 diabetes^2^ at genomewide significance. Examination of regions associated with both diseases could lead to uncovering signals that simultaneously alter disease risk for both diseases, termed colocalisation. Uncovering colocalising signals could provide biological insights into shared disease mechanisms, and potentially reveal therapeutic targets effective for both diseases. A recent analysis suggested that the same genetic variant is altering risk of both type 1 and type 2 diabetes in five regions, near Centromere Protein W (*CENPW*), Chymotripsinogen B1 (*CTRB1*)/Breast Cancer Anti-Estrogen Resistance Protein 1 (*BCAR1*), GLIS Family Zinc Finger Protein 3 (*GLIS3*), B-cell Lymphoma 11A (*BCL11A*) and Thyroid Adenoma-Associated Protein (*THADA*)^3^.

Here, we identified all regions across the genome that showed evidence of association to both type 1 and type 2 diseases at false discovery rate (FDR) <0.01, and assessed colocalisation between the two diseases in each of these regions. Furthermore, to account for the possibility of multiple causal variants within an associated region, we extended the analysis to investigate conditionally-independent associations within each region, to assess whether any of the associations with one disease colocalised with any associations in the other.

## Methods

Type 1 diabetes meta-analysis summary statistics were generated using genome wide association study (GWAS) data from 3,983 cases and 3,994 controls from the UK genotyped using the Illumina Infinium 550K platform, 1,926 cases and 3,342 controls from the UK genotyped using the Affymetrix GeneChip 500K platform and 1,558 cases and 2,882 controls from Sardinia genotyped using the Affymetrix 6.0 and Illumina Omni Express platforms, totalling 7,467 cases and 10,218 controls (**ESM Table 1**). Genotypes were imputed using the Haplotype Reference Consortium (HRC) reference panel for the UK collections ^4^, and a custom Sardinian reference panel of 3,514 Sardinians for the Sardinian collection (**ESM - Imputation**).

Summary statistics for type 2 diabetes were from 74,124 cases and 824,006 controls of European ancestry, imputed using the HRC reference panel^2^. Regions associated with both diseases were identified by selecting all variants with type 1 diabetes and a type 2 diabetes association with FDR<0.01(**ESM – Type 1 diabetes GWAS**). In each such region, windows of approximately 0.5 Mb were taken to examine colocalisation (**ESM - Regions associated with both diseases**). Within these regions, forward stepwise logistic regressions were carried out for both diseases, and conditional summary statistics were obtained so each conditionally-independent signal from both diseases could be tested against each other for colocalisation (**ESM - Conditional analyses**).

Colocalisation of signals was assessed using *coloc*^5^, a Bayesian method that enumerates the posterior probability that the association signals in a region are shared between traits. The prior probability of association with either disease was taken to be 1×10^−4^ and the prior probability that the association signal is shared across traits was taken to be 5 × 10^−6^, as recommended^6^. The threshold to consider signals as colocalising was conservatively chosen at a posterior probability ≥0.9. Colocalisation was also examined using an alternative approach, as a secondary analysis, eCAVIAR ^7^ (**ESM – eCAVIAR**).

## Results

Including conditionally-independent association signals, 81 colocalisation analyses were carried out across 42 chromosomal regions that showed association to both diseases (**ESM Table 2**).

Four signals showed evidence of colocalisation using *coloc*, and they were also the regions with the highest *eCAVIAR* regional colocalisation posterior probabilities (**ESM Table 3**). The first was on chromosome 16q23.1, near *CTRB1* and *BCAR1*, with a posterior probability of colocalisation (H4PP, hereafter) of 0.98 (**ESM Figure 1**). The minor A allele at the type 2 diabetes index variant, rs72802342 (C>A), is protective for type 2 diabetes (OR=0.87, p=4.00×10^−32^) and susceptible for type 1 diabetes (OR=1.33, p=5.81×10^−10^).

The second was on chromosome 11p15.5, near *INS*, where the primary type 2 diabetes association colocalised with the secondary type 1 diabetes association (H4PP=0.95, **ESM Figure** 2). The direction of effect was opposite, with the minor A allele at the type 2 diabetes index variant, rs4929965 (G>A), associated with susceptibility to type 2 diabetes (OR=1.07, p=4.8O×10^−25^) and protection from type 1 diabetes (OR=0.87, p=1.89×10^−5^).

Thirdly, a region on chromosome 4p16.3 colocalised (H4PP=0.97) (**Figure 1),** near Transmembrane protein 129 (*TMEM129*). The minor T allele at the type 2 diabetes index variant, rs56337234 (C>T), was associated with decreased risk of type 2 diabetes (OR=0.94, p=1.4×10^−17^) and increased risk of type 1 diabetes (OR=1.12, p=4.07×10^−6^).

**Figure 1:**
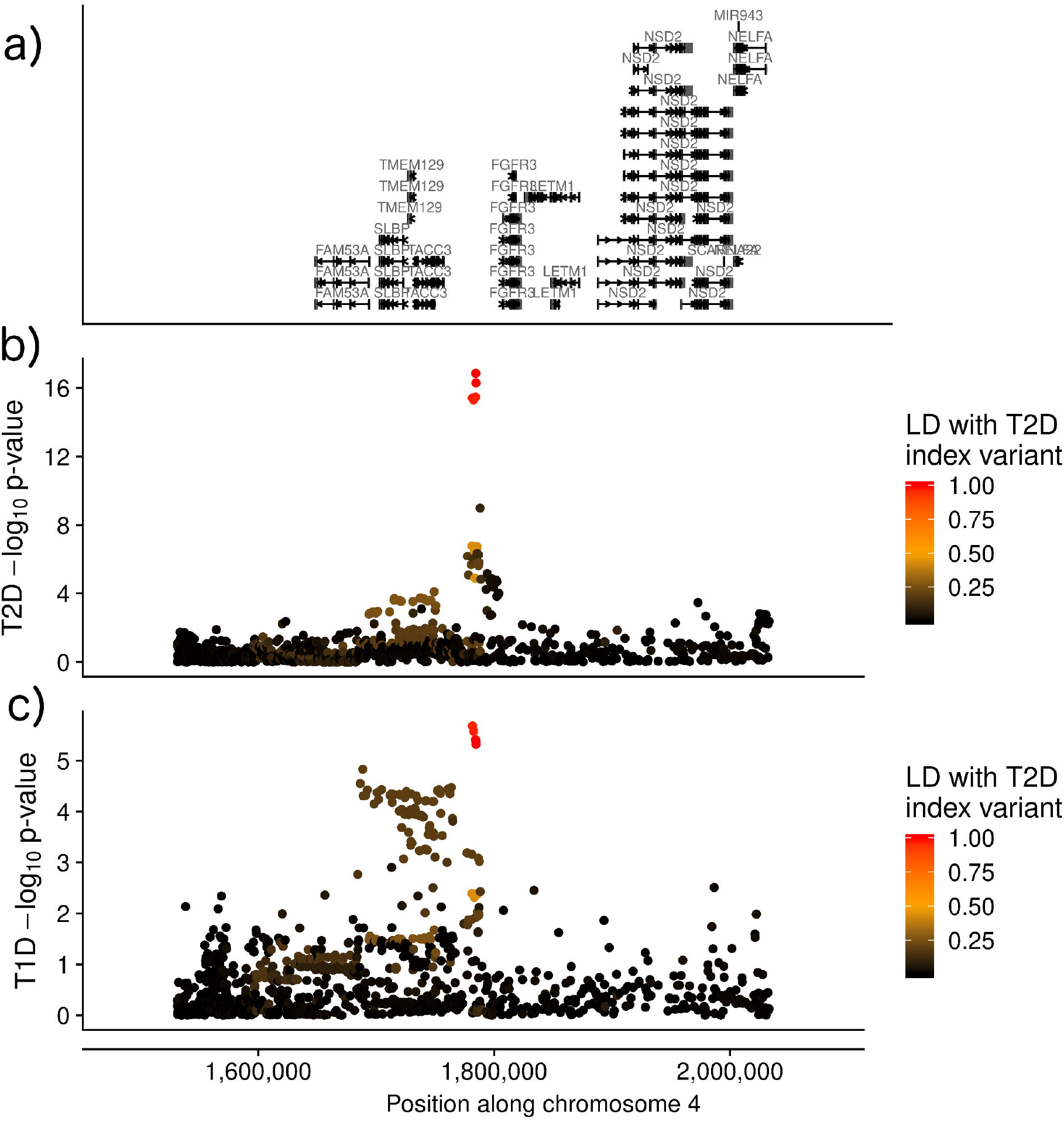
Manhattan plots showing a) gene locations and −log_10_p-value of association for each variant by position along chromosome 4 (genome build 37) in the *TMEM129* region for b) type 2 diabetes and c) type 1 diabetes, coloured by r^2^ to the type 2 diabetes index variant, rs56337234.

Finally, a region on chromosome 1p31.3, near Phosphoglucomutase 1 (*PGM1*), colocalised (H4PP=0.91, **ESM Figure 3),** with the minor T allele at the type 2 diabetes index variant rs2269247 (C>T) decreasing risk of type 2 diabetes (OR=0.96, p=4.6×10^−7^) and increasing risk of type 1 diabetes (OR=1.15, p=1.9×10^−6^) (**Table 1**).

**Table 1:**
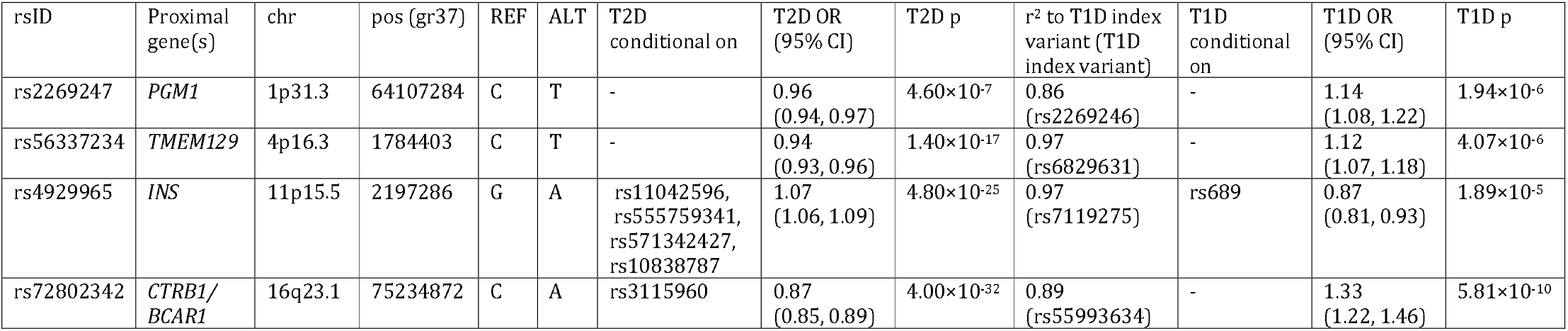
Regions with a colocalisation posterior probability of ≥0.9 between type 1 diabetes and type 2 diabetes. Summary statistics given from the perspective of the index type 2 diabetes variant and with respect to the ALT allele. T2D= type 2 diabetes, T1D= type 1 diabetes, r^2^ obtained from 1000 Genomes Project European population.

| rsID | Proximal gene(s) | chr | pos (gr37) | REF | ALT | T2D conditional on | T2D OR (95% CI) | T2D p | $r^2$ to T1D index variant (T1D index variant) | T1D conditional on | T1D OR (95% CI) | T1D p |
| --- | --- | --- | --- | --- | --- | --- | --- | --- | --- | --- | --- | --- |
| rs2269247 | *PGM1* | 1p31.3 | 64107284 | C | T | - | 0.96 (0.94, 0.97) | $4.60 \times 10^{-7}$ | 0.86 (rs2269246) | - | 1.14 (1.08, 1.22) | $1.94 \times 10^{-6}$ |
| rs56337234 | *TMEM129* | 4p16.3 | 1784403 | C | T | - | 0.94 (0.93, 0.96) | $1.40 \times 10^{-17}$ | 0.97 (rs6829631) | - | 1.12 (1.07, 1.18) | $4.07 \times 10^{-6}$ |
| rs4929965 | *INS* | 11p15.5 | 2197286 | G | A | rs11042596, rs555759341, rs571342427, rs10838787 | 1.07 (1.06, 1.09) | $4.80 \times 10^{-25}$ | 0.97 (rs7119275) | rs689 | 0.87 (0.81, 0.93) | $1.89 \times 10^{-5}$ |
| rs72802342 | *CTRB1/BCAR1* | 16q23.1 | 75234872 | C | A | rs3115960 | 0.87 (0.85, 0.89) | $4.00 \times 10^{-32}$ | 0.89 (rs55993634) | - | 1.33 (1.22, 1.46) | $5.81 \times 10^{-10}$ |

We did not replicate the finding that the chromosome regions near *CENPW, GLIS3, BCL11A or THADA* colocalised between type 1 and type 2 diabetes (H4PP *CENPW*=0.12, *GLIS3*=0.29, *BCL11A*=0.28, *THADA* not examined as no type 1 diabetes association existed in the region (FDR=0.07)). To investigate these discrepancies, we examined two other large type 2 diabetes meta-analyses: a trans-ethnic study including 1,407,282 individuals ^8^ and a study of 433,540 individuals of East Asian ancestry^9^. For the *CENPW* and *BCL11A* regions, the type 2 diabetes signal is consistent with at least one of the other GWAS studies (measured by linkage disequilibrium (LD) in Europeans to the other study index variants, **ESM Table 4);** and, the type 1 diabetes index variant is not in strong LD (r^2^<0.41) with any of the index variants for type 2 diabetes across the three GWAS studies. However, at *GLIS3*, there appears to be a distinct signal in the European study^2^ compared to the trans-ethnic and East Asian type 2 diabetes studies (r^2^=0.65), and the index variants from these two studies are in higher r^2^ with the type 1 diabetes signal in our analysis (r^2^=0.68), and even higher r^2^ with the index variant from a larger T1D genetic analysis^1^ (r^2^=0.99), indicating that the signal near *GLIS3* does colocalise between type 1 and type 2 diabetes with concordant direction of effect, as previously identified^10^.

## Discussion

Using genetic association summary statistics from European populations, we identified 42 regions that showed association with both type 1 and type 2 diabetes, with 81 conditionally-independent association signals across those regions. Four signals (near *CTRB1/BCAR1, INS, TMEM129* and *PGM1*) colocalised between the diseases, including a signal at the complex *INS* region for the first time, which was achieved by examining conditional summary statistics. However, in all four cases, the allele increasing risk for one disease was protective against the other. Examination of additional trans-ethnic and East Asian type 2 diabetes genetic analyses, indicated that a fifth association, near *GLIS3*, is likely to colocalise between diseases, with concordant direction of effect.

Given the distinct mechanisms underlying β-cell dysfunction and cell death between the two diseases ^11^, it is perhaps unsurprising that no additional signals were detected with concordant direction of effect. However, the type 1 diabetes GWAS was much smaller than the type 2 diabetes analysis, and therefore had less statistical power to detect more subtle genetic effects. If a type 1 diabetes GWAS were to be performed with similar power to the type 2 diabetes GWAS, more regions might colocalise between the two diseases, but either the effects of these additional regions on type 1 diabetes would be small compared to the currently known associations, or they would be rare variants with larger effect sizes.

That four of five colocalisation signals had opposite directions of effect implies a complex genetic relationship between the two diseases. Whilst the directional discordance offers little hope for effective treatments for both diseases simultaneously at these particular targets, it can offer biological insight into the disease pathways that these regions act upon, and even if there is directional discordance, the genetics could be highlighting the same therapeutic target. We did not replicate the findings that the associations near *BCL11A, CENPW* and *THADA* colocalise between the two diseases^3^, despite overlapping samples and similar numbers of cases and controls in the type 1 diabetes GWAS. This is likely due to three reasons: i) the previous study^3^ examined colocalisation using weaker association signals, for example, the colocalisation near *THADA* was based on a type 1 diabetes association p-value of 0.01; ii) we used a more stringent prior for colocalisation between the two diseases, as recently suggested ^6^ (5×10^−6^ vs. 1×10^−5^); and iii) we used a more stringent posterior probability threshold to declare colocalisation (0.9 vs. 0.5). Our increased stringency compared to the previous analysis ^3^, whilst increasing the probability that any identified shared signals will be true positives, may have decreased our sensitivity to detect all colocalisations. For example, by examining other large type 2 diabetes GWAS analyses and a larger type 1 diabetes genetic analysis, we conclude that the association near *GLIS3* likely does colocalise between the two diseases, and with concordant directions of effect.

In conclusion, with current GWAS sample sizes, just five associations appear to colocalise between type 1 diabetes and type 2 diabetes, four with opposing direction of effect. Larger sample sizes would be required to identify the depth of genetically identified therapeutic targets to treat or prevent both diseases simultaneously.

## Supporting information

ESM

## Acknowledgments

We gratefully acknowledge all participants for allowing the analysis of their genotypic and phenotypic data.

## Data availability

Type 1 diabetes summary statistics will be available through GWAS catalog (https://www.ebi.ac.uk/gwas/). Type 2 diabetes summary statistics are already publicly available.

## Authors’ relationships and activities

Mark McCarthy has served on advisory panels for Pfizer, Novo Nordisk and Zoe Global, has received honoraria from Merck, Pfizer, Novo Nordisk and Eli Lilly, and research funding from Abbvie, Astra Zeneca, Boehringer Ingelheim, Eli Lilly, Janssen, Merck, Novo Nordisk, Pfizer, Roche, Sanofi Aventis, Servier, and Takeda. As of June 2019, Mark McCarthy is an employee of Genentech, and a holder of Roche stock. Anubha Mahajan is an employee of Genentech since January 2020, and a holder of Roche stock.

Jamie Inshaw is an employee of Exploristics since June 2020.

John Todd serves on the advisory board of GSK.

## Contribution statement

JRJI carried out the type 1 diabetes meta-analysis and the colocalisation analyses, drafted the manuscript and approved the final version.

AM carried out the type 2 diabetes meta-analysis and conditional analyses, revised the article for intellectual content and approved the final version.

CS and FC were involved in data collection in the Sardinia collection and carried out the association testing in this collection, revised the article for intellectual content and approved the final version.

DC provided statistical advice and input, and made contributions to interpretation of the data, revised the article for intellectual content and approved the final version..

MIS provided biological insight, contributed towards interpretation of the data, revised the article for intellectual content and approved the final version.

MM and JAT oversaw the research, contributed towards the conception, design and data collection, revised the article for intellectual content and approved the final version.

JAT is the guarantor of this work.

