## Supplementary material for "Analysis of overlapping genetic association in type 1 and type 2 diabetes": ESM

**Electronic supplementary material**

**Methods**

***Imputation***

Imputation for the type 1 diabetes analysis was performed using the Michigan Imputation server, pre phasing using SHAPEIT2 and imputation using Minimac3 ^1^. Prior to imputation, variants out of Hardy-Weinberg Equilibrium (p-value<1×10^-6^), rare variants (minor allele frequency<0.01) and variants with high missing call rate (>0.95) were excluded. Remaining variants were then aligned to the HRC reference panel strand using the following pipeline https://www.well.ox.ac.uk/~wrayner/tools/.

***Type 1 diabetes GWAS***

Association of each variant with type 1 diabetes was assessed using *SNPTEST*, the ‘newml’ method ^2^, adjusting for the three largest principal components within that collection from a pruned (r^2^<0.2) genetic matrix without rare variants (MAF>0.01).

The UK collections were combined in an inverse-variance weighted meta-analysis. However, prior to meta-analysis, variants were excluded from the results in that collections if:

1) a variant had an imputation information score of <0.3 in cases or controls

2) the difference in imputation information score between cases and controls was >0.05

Following the UK meta-analysis, variants were excluded based on the following criteria:

1) if there was a difference in MAF in controls between the Affymetrix and the Illumina collections of >0.05

2) if there was a difference in MAF in cases between the Affymetrix and the Illumina collections of >0.05

3) if there was a difference in MAF of >0.05 between controls and the HRC reference panel MAF

4) if the difference in log-odds ratio estimate between the Affymetrix and Illumina collections was >0.5

Once the UK estimates were obtained, the UK-Sardinia meta-analysis was carried out, including only variants with MAF>0.01 in Sardinians into the meta-analysis (those with MAF<0.01 in Sardinians but included in the UK analysis would be included in the final results but with only the UK results contributing towards the overall association statistic). Any variant excluded in the UK-ancestry analysis was also excluded from the UK-Sardinia meta-analysis.

***Regions associated with both diseases***

To identify regions to examine in colocalisation analyses, we first calculated the false discovery rate (FDR) value for each variant after excluding the HLA region in the type 1 diabetes analysis. Once an associated region was identified from the set of genome wide associations, a 0.5Mb window around the index variant was excluded and placed in the list of regions for downstream analyses. Then the next most associated variant was identified and a 0.5Mb region around this variant was added to this list of regions for downstream analysis. This process was repeated until no variants were left with an FDR<0.01. This process identified 98 type 1 diabetes 0.5Mb regions for downstream analysis.

The same process was then performed for type 2 diabetes, without exclusion of the HLA region to calculate the FDR value for each variant. This process identified 852 type 2 diabetes 0.5Mb regions for downstream analysis.

All overlapping regions were then kept for conditional analyses, and colocalisation analyses, taking the union of the overlapping regions as the region to analyse.

***Conditional analyses***

Forward stepwise conditional regression for type 1 diabetes was carried out using UK data only, performed using the Affymetrix and then the Illumina data, before meta-analysing. The procedure was stopped when a variant added to the model had a Wald test meta-analysis p value of >6.25×10^-6^, which was the maximum p-value from univariable analyses with a false discovery rate (FDR)<0.01. Once all conditionally independent associations were identified, then all conditionally independent signals were included in the model to re-examine the association in the primary association signal.

Forward stepwise conditional regression for type 2 diabetes was carried out using the ‘cojo’ option in GCTA ^3^.

***eCAVIAR***

*eCAVIAR* analyses ^4^ were performed using the T1D Illumina cohort to generate an LD matrix for variants included in the analysis, and this structure was assumed to be consistent for the T1D and T2D datasets. The same variants were included in the analyses as in the *coloc* analysis. We performed analyses in the same way as in the *coloc* analysis, by conditioning on other association signals in the region and examining colocalisation using conditional summary statistics where relevant. We therefore assumed the maximum number of causal variants for each colocalisation analysis was 1.

eCAVIAR enumerates the colocalisation posterior probability (CLPP) for each variant included in the analysis. In order to obtain an estimate for colocalising signals across the region, which is more similar to the hypothesis *coloc* is testing, we summed each variant CLPP to obtain the eCAVIAR regional CLPP, which are reported in **ESM Table 3**.

***Code availability***

Code used to carry out this analysis is available at https://github.com/jinshaw16/t1d-t2d-colocalisation.

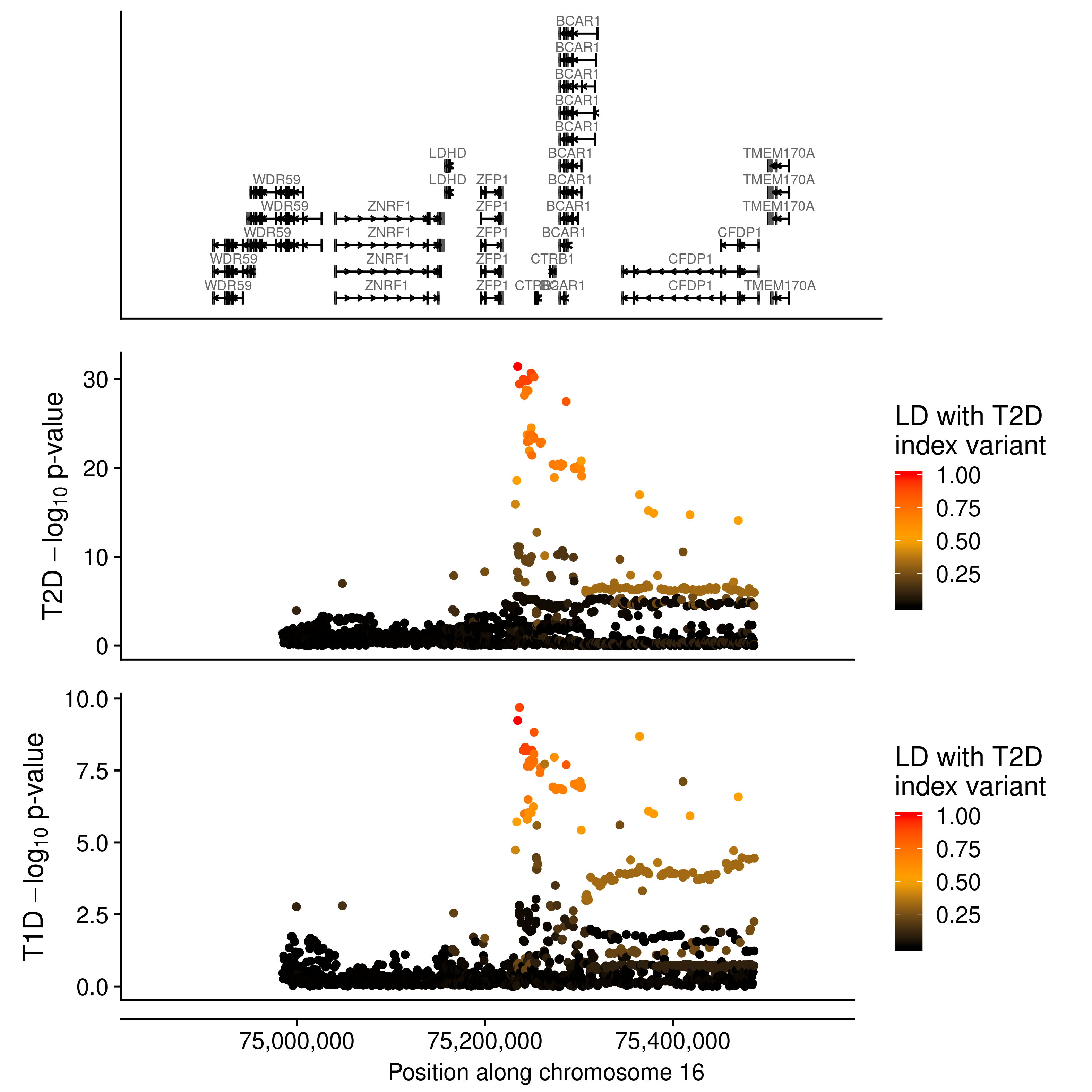

**ESM Figure 1:** Manhattan plots showing –log_10_p-value of association for each variant by position along chromosome 16 (genome build 37) in the *CTRB1/BCAR1* region for type 2 diabetes (middle panel) and type 1 diabetes (bottom panel), coloured by r^2^ to the type 2 diabetes index variant, rs72802342.

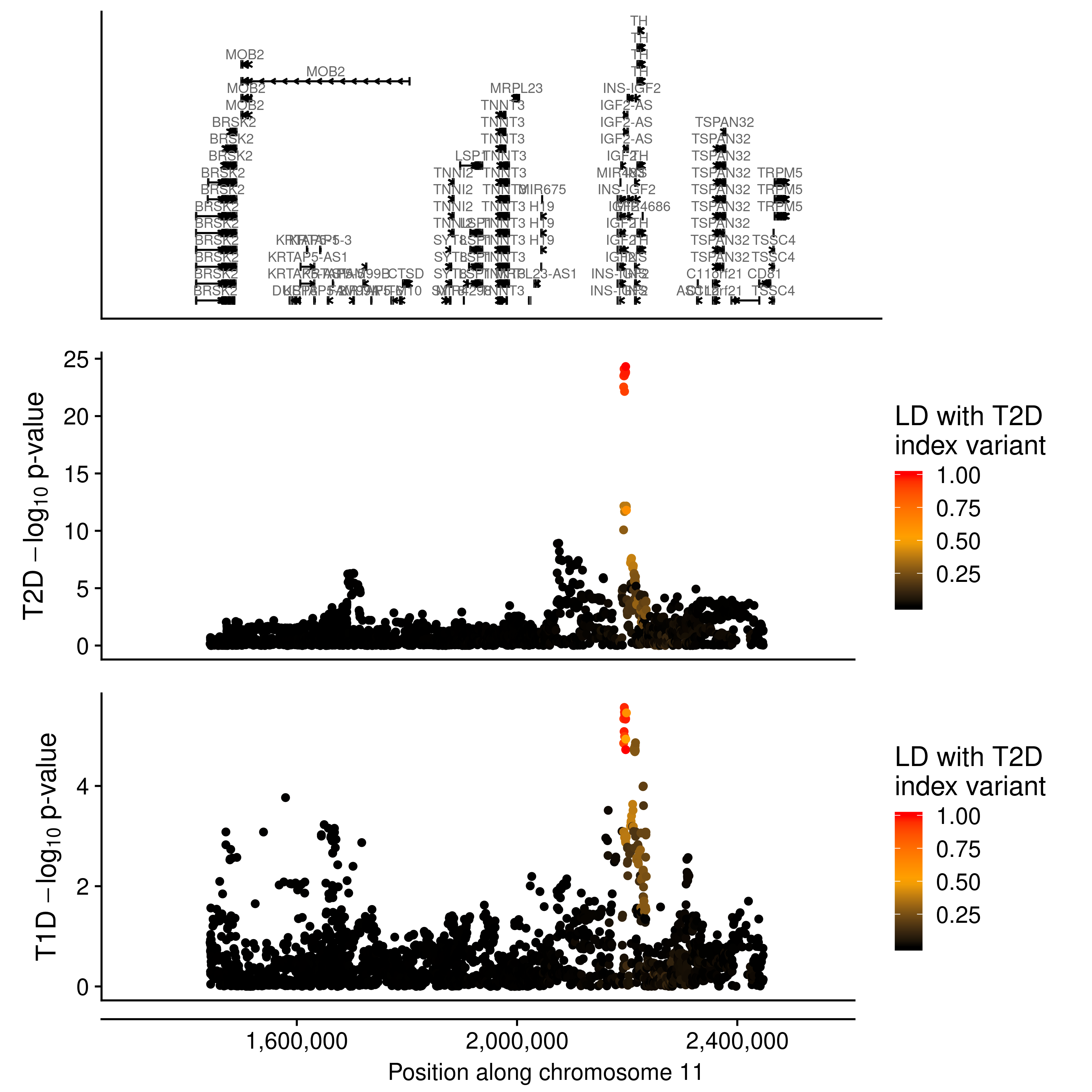

**ESM Figure 2:** Manhattan plots showing –log_10_p-value of association for each variant by position along chromosome 11 (genome build 37) in the *INS* region for type 2 diabetes (middle panel) and type 1 diabetes (bottom panel) , conditional on primary signal index variant rs689, coloured by r^2^ to the type 2 diabetes index variant, rs4929965.

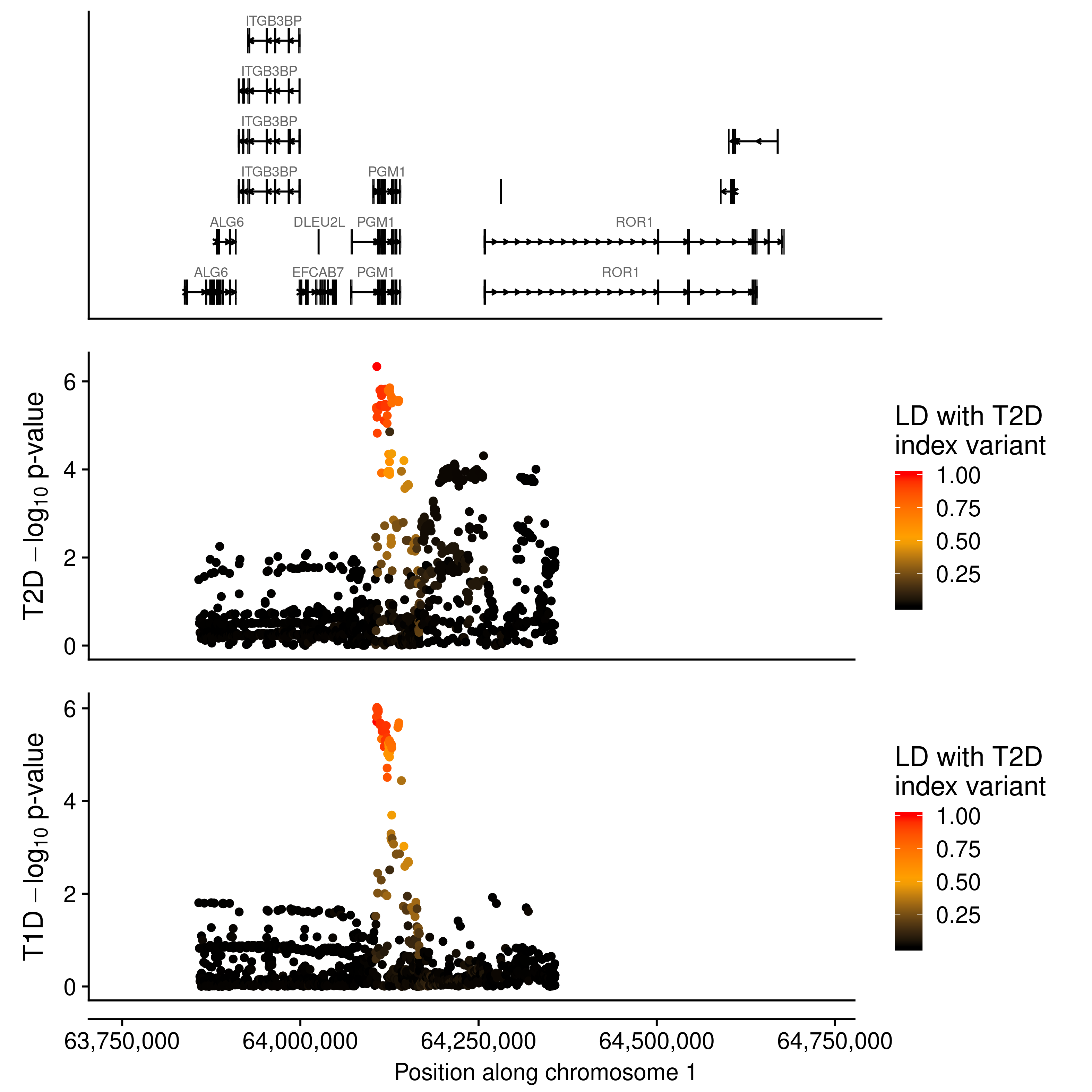

**ESM Figure 3:** Manhattan plots showing –log_10_p-value of association for each variant by position along chromosome 1 (genome build 37) in the *PGM1* region for type 2 diabetes (middle panel) and type 1 diabetes (bottom panel), coloured by r^2^ to the type 2 diabetes index variant, rs2269247.
